# Embryonic depletion of D-aspartate perturbs NMDA receptor-dependent long-term potentiation in the hippocampus of juvenile mice

**DOI:** 10.64898/2026.04.22.720120

**Authors:** Dalila Mango, Francesco Errico, Zoraide Motta, Sepideh Dashtiani, Anna Di Maio, Robert Nisticò, Maria Egle De Stefano, Loredano Pollegioni, Alessandro Usiello

**Affiliations:** School of Pharmacy, Department of Biology, University of Rome ‘Tor Vergata’, Via della Ricerca Scientifica, 00133 Rome, Italy; CEINGE Biotecnologie Avanzate “Franco Salvatore”, Naples, Italy; Department of Agricultural Sciences, University of Naples “Federico II”, Portici, Italy; “The Protein Factory 2.0”, Dipartimento di Biotecnologie e Scienze della Vita, Università degli Studi dell’Insubria, Varese, Italy; Department of Physiology and Pharmacology “V. Erspamer”, Sapienza University of Rome, Rome, Italy; Department of Environmental, Biological and Pharmaceutical Sciences and Technologies, Università degli Studi della Campania “Luigi Vanvitelli”, Caserta, Italy; Department of Biology and Biotechnologies “Charles Darwin”, Sapienza University of Rome, Rome, Italy; Center for Research in Neurobiology “Daniel Bovet”, Sapienza University of Rome, Rome, Italy

**Keywords:** D-amino acids, neurodevelopment, neuropsychiatric disorders, synapse, sex

## Abstract

D-Aspartate (D-Asp) is an endogenous D-amino acid that exhibits a pronounced developmental peak in the mammalian brain, suggesting a potential regulatory role in glutamatergic signaling and neurodevelopment. Disruption of D-Asp homeostasis has been associated with neuropsychiatric disorders characterized by early-life circuit vulnerability, including schizophrenia and autism spectrum disorders. However, its functional impact to hippocampal physiology remains incompletely defined. Here, we investigated how constitutive D-Asp depletion affects synaptic function in the hippocampal CA1 region of *Ddo*-knock-in (*Ddo*-KI) mice, in which zygotic overexpression of the D-Asp-degrading enzyme, D-aspartate oxidase (DASPO), results in embryonic and persistent D-Asp deficiency. Electrophysiological recordings were performed in acute hippocampal slices from male and female mice at postnatal day 30 (P30) and day 60 (P60). Basal synaptic transmission, assessed through paired-pulse ratio and spontaneous excitatory/inhibitory events, was unaltered between genotypes, indicating preserved presynaptic release probability and overall excitation/inhibition balance. In contrast, NMDA receptor (NMDAR)-dependent synaptic plasticity was selectively altered, as theta-burst stimulation induced significantly greater long-term potentiation (LTP) in juvenile P30 *Ddo*-KI mice, whereas this difference was no longer observed at P60. Consistently, patch-clamp recordings revealed a reduced AMPAR/NMDAR ratio in P30 *Ddo*-KI males, suggesting an increased relative contribution of NMDAR-mediated currents. Importantly, acute bath application of exogenous D-Asp restored LTP to wild-type levels, demonstrating rapid reversibility and supporting a model of homeostatic receptor rebalancing rather than irreversible circuit alterations. Biochemical assays confirmed significantly increased DASPO activity and reduced D-Asp levels in *Ddo*-KI mice. However, these parameters remained stable between P30 and P60, indicating that the age-dependent plasticity phenotype is unlikely to arise from progressive biochemical changes. Together, these findings indicate that developmental D-Asp deficiency induces a transient, juvenile-specific alteration characterized by enhanced NMDAR-dependent synaptic plasticity, which can be rapidly normalized upon D-Asp re-exposure.

## 1. Introduction

D-aspartate (D-Asp) is an atypical endogenous D-amino acid that has emerged as a key signaling molecule in the mammalian brain and neuroendocrine system. Unlike its canonical L-enantiomer, L-aspartate, D-Asp is not incorporated into proteins but exists in free form and participates in distinct neuronal and endocrine regulatory pathways (D’Aniello, 2007; Ota et al., 2012; Errico et al., J Pharm Biomed Anal, 2015; Errico et al, BBA, 2020; Usiello et al, Int J Mol Sci, 2020). Pharmacologically, D-Asp acts as an endogenous agonist of N-methyl-D-aspartate receptors (NMDARs), binding to the L-glutamate site on the GluN2 subunit and modulating excitatory glutamatergic neurotransmission (Krashia et al., 2016; reviewed in Souza et al., 2023; Errico et al., 2015 and 2020). Consistent with this role, electrophysiological studies have shown that exogenous D-Asp evokes NMDAR-dependent inward currents in neurons and a transient rise in intracellular Ca^2^□ levels, which are abolished by both competitive (D-AP5) and non-competitive (MK-801) NMDAR antagonists (Errico et al., 2008a,b; Errico et al., 2011a). In addition to NMDAR ionotropic signaling, D-Asp also influences metabotropic glutamate receptor 5 (mGluR5) activity (Molinaro et al., 2010), further contributing to intracellular pathways relevant to synaptic plasticity.

Importantly, increased D-Asp availability has been shown to enhance glutamatergic transmission *in vivo*. In particular, elevated D-Asp levels in *Ddo* knockout mice, lacking the D-Asp-degrading enzyme D-aspartate oxidase (DASPO), or following chronic D-Asp administration, enhance NMDAR-dependent long-term potentiation (LTP) in the hippocampus and improve performance in hippocampal-dependent cognitive tasks (Errico et al., 2008a,b; Errico et al., 2011a,b; Errico et al., 2014). These findings support a facilitatory role of D-Asp in synaptic plasticity and memory processes.

In addition to postsynaptic effects, *in vivo* microdialysis in freely moving mice and synaptosome studies from the prefrontal cortex (PFC) have shown that D-Asp enhances cortical presynaptic L-glutamate release via stimulation of AMPA, NMDA, and mGlu receptors located on nerve terminals (Sacchi et al., 2017). These findings support a facilitatory role for both ionotropic and metabotropic auto-receptors in presynaptic modulation, at least within this brain region.

A distinctive feature of D-Asp in the central nervous system is its marked developmental regulation. Brain levels peak during prenatal and early postnatal development and decline sharply after birth (Dunlop et al., 1986; Hashimoto et al. 1993; Hashimoto et al.,1995; Neidle and Dunlop, 1990; Sakai et al., 1998; Wolosker et al., 2000; Punzo et al., 2016; De Rosa et al., 2020), due to the postnatal activation of the degradative enzyme DASPO, encoded by the *Ddo* gene (Van Veldhoven et al., 1991; Katane and Homma, 2010; Molla et al., 2020; Punzo et al., 2016; De Rosa et al., 2020). This tightly controlled temporal profile suggests that D-Asp plays a critical role during brain development, particularly in processes related to circuit formation and maturation.

Accordingly, dysregulation of D-Asp metabolism has been reported in clinical and preclinical studies and implicated in neurodevelopmental and psychiatric disorders characterized by glutamatergic dysfunction, including schizophrenia and autism spectrum disorders (ASD) (Errico et al., 2013; Errico et al., 2015; Nuzzo et al., 2017; De Rosa et al., 2022; Lombardo et al., 2022; et al., 2024; Rampino et al., 2024; Di Maio et al., 2025), as well as during developmental stages in neuromuscular disorders, such as Duchenne Muscular Dystrophy (Mastrostefano et al., 2025).

To directly investigate the consequences of developmental D-Asp dysregulation, a genetically engineered knock-in (KI) mouse model (R26^*Ddo/Ddo*^) was generated, in which DASPO overexpression is driven from the zygotic stage, resulting in constitutive D-Asp depletion from embryogenesis through adulthood (De Rosa et al., 2020). This mouse model shows abnormally enhanced memory alongside impaired sociability, phenotypes accompanied by reduced parvalbumin-positive interneurons in the PFC (De Rosa et al., 2020; Lombardo et al 2022). At the molecular level, neonatal brain metabolomic and proteomic analysis revealed significant alterations in pathways related to GABAergic and glutamatergic neurotransmission, as well as brain development and cytoskeletal organization (Grimaldi et al., 2020; Errico et al., 2025), indicating that D-Asp exerts broad neurodevelopmental effects beyond synaptic modulation. Additionally, developmental D-Asp depletion leads to reduced neurogenesis in the developing cortex and to decreased cortical and striatal volume in adulthood (Lombardo et al., 2022).

In the present study, we investigated how early and sustained D-Asp depletion affects hippocampal basal synaptic transmission, NMDAR-dependent LTP, and AMPAR/NMDAR current ratios in juvenile (P30) and adult (P60) *Ddo*-KI mice, analyzing males and females separately.

## 2. Material and methods

### 2.1 Animals

Both male and female *Ddo*-KI mice, together with their corresponding wild-type (WT) controls, were examined at postnatal day 30 (P30) (male, n = 29; female, n = 23) and 60 (P60) (male, n = 9; female, n = 5). These two ages were selected in accordance with previous studies (De Rosa et al., 2022), as they correspond to the juvenile (P30) and young adult (P60) stages, a postnatal developmental window in which hippocampal maturation is substantially established. No sample size was calculated a priori, but the number of samples used for the study was determined based on our previous studies (Cristino et al 2015; De Jaco et al., 2017).

All procedures were conducted in accordance with national regulations (D.Lgs 26/ 2014), international guidelines for animal welfare (European Communities Council Directive 2010/63/EU), and were approved by the EBRI Rita Levi-Montalcini Foundation (F8BBD.N.I6P). Every effort was made to reduce the number of animals used and to minimize discomfort, in accordance with the 3Rs principle. *Ddo*-KI mice were generated in-house as previously described (De Rosa et al., 2020) and all mice were housed in groups (n = 4-5) per standard cage (29 × 17.5 × 12.5 cm) under controlled temperature (22 ± 1 °C) and a 12 h-hour light/dark cycle, with food and water available ad libitum (Plaisant S.r.l.). For tissue collection, mice were rapidly decapitated without prior anesthesia, in accordance with approved protocols, to preserve tissue viability for electrophysiological and biochemical analyses.

All electrophysiological recordings, HPLC detections and enzymatic activity assays were blinded, and the experimental group assignment was revealed during data analysis.

### 2.2 Extracellular field recordings

Parasagittal hippocampal slices (350 μm thick) were prepared from male and female *Ddo*-KI mice and their corresponding WT controls at P30 (male, n=20; female=18) and P60 (male, n = 18; female, n = 10), following established procedures (Mango et al., 2017). Field excitatory post-synaptic potential (fEPSP) was evoked using a stimulating electrode placed near the Schaffer collateral fibers and recorded with an electrode placed in the CA1 *stratum radiatum*. Short-term plasticity was assessed by measuring the paired pulse ratio (PPR) at inter-pulse intervals of 50, 100, 200, 300, 400, and 500 ms, calculated as the ratio of the second to the first fEPSP amplitude. LTP was elicited using a theta burst stimulation (TBS) protocol (Piccioni et al., 2024).

### 2.3 D-Asp perfusion

To rescue synaptic plasticity, hippocampal sections from P30 male *Ddo*-KI mice were acutely perfused with 2 μM D-Asp, dissolved in cerebrospinal fluid (CSF), 30 min before TBS. Potentiation magnitude was then recorded 60 min after the conditioning stimulus.

### 2.4 Whole-cell patch clamp recordings

Whole-cell patch-clamp recordings were obtained from hippocampal CA1pyramidal neurons according to standard protocols (Mango et al., 2017; De Jaco et al., 2017). As above, parasagittal hippocampal slices (250 μm thick) were prepared from P30 male and female *Ddo*-KI mice and their corresponding WT. For each neuron, spontaneous excitatory postsynaptic currents (sEPSC) were recorded at the reversal potential of the GABAA receptors (-70 mV), whereas spontaneous inhibitory postsynaptic currents (sIPSC) were recorded at the reversal potential for ionotropic glutamate receptors (+10 mV). The excitatory/inhibitory (E/I) balance was calculated as the ratio of sEPSC to sIPSC frequency (De Jaco et al., 2017).

For paired-pulse (PP) experiments (50 ms inter-pulse interval), excitatory postsynaptic currents were evoked via a stimulating electrode placed in the CA1 *stratum radiatum*. The paired-pulse ratio (PPR) was calculated as the amplitude ratio of the second to the first EPSC.

The AMPAR/NMDAR ratio was quantified as the ratio between the peak EPSC amplitude at -80 mV holding potential (AMPAR component), and the peak EPSC amplitude at +40 mV holding potential (NMDAR component) over a 2 ms window beginning 60 ms after the stimulation artifact, as described previously (De Jaco et al., 2017).

### 2.5 DASPO activity assay

Hippocampal samples from *Ddo*-KI mice and their corresponding WT controls at P30 (WT, n = 4; *Ddo*-KI, n=5) and P60 (WT, n = 4; *Ddo*-KI, n=4) were resuspended and homogenized using a pellet micro-pestle in 20 mM sodium phosphate buffer (pH 8.0) containing 0.1% Triton X-100 (93426, Fluka), supplemented with complete protease inhibitor cocktail (11836153001, Roche) and phosphatase inhibitor cocktail (5870, Cell Signaling Technology). Samples were prepared at a ratio of 1:10 or 1:20 (w/v), depending on tissue availability. Following homogenization, lysates were sonicated (10 s x 3 cycles) and centrifuged at 16,000 g for 20 min at 4 °C. H□O□ production from DASPO reaction was measured using an Amplex UltraRed-based assay (Rosini et al., 2018). Tissue extracts (5–15 μL) were added to a reaction mixture containing 20 μM Amplex UltraRed (Thermo Fisher Scientific), 0.05 U/mL horseradish peroxidase, 2.5 mM NaN□, 5 μM FAD, and 80 mM D-Asp. Fluorescence (Ex = 535 nm, Em = 590 nm) was recorded at 30, 60, 120, 240, and 360 min, and overnight at 25 °C using a Tecan Infinite M Plex microplate reader. A calibration curve was generated with 0.1–10 μM H□O□. Control reactions with recombinant DASPO (0.04 mU), without tissue extract or substrate, and adding 20 mM meso-tartaric acid (a DASPO inhibitor) were performed simultaneously. Fluorescence values measured in the presence of the inhibitor were subtracted to eliminate nonspecific signal. Protein concentration was determined by Bradford assay. Data are reported as µU/µg protein at 360 min (time of maximal activity).

### 2.7 Enantiomeric HPLC

Hippocampal samples from *Ddo*-KI mice and their corresponding WT controls at P30 (WT, n = 2; *Ddo*-KI, n=3) and P60 (WT, n = 4; *Ddo*-KI, n=4) were analyzed following the protocol reported in (Punzo et al., 2016), with some methodological adaptations. Samples were resuspended in ice-cold 0.2 M trichloroacetic acid (TCA) at a ratio of 1:10 or 1:20 (w/v), depending on tissue availability. Homogenization was performed using a manual pellet micro-pestle, followed by sonication (10 s x 3 cycles) and centrifugation at 16,000 g for 20 min at 4 °C. The precipitated protein pellets were then resuspended in 1% SDS, and total protein concentration was measured by Bradford assay. For amino acid analysis, 5 μL of the TCA extracts were neutralized with NaOH and derivatized before injection with o-phthaldialdehyde (OPA) and N-acetyl-L-cysteine (NAC) in 50% methanol. Separation of diastereomeric derivatives was performed under isocratic conditions on a C8 reversed-phase column (Symmetry, 3.5 μm, 4.6 × 150 mm; Waters). 0.1 M sodium acetate buffer (pH 6.2) containing 1% tetrahydrofuran (THF) was used as mobile phase, at 0.7–1.0 mL/min flow rate. All amino acids analysed were identified and quantified by comparison of retention times and peak areas with those of external pure standards. D-Asp and D-Ser levels were further confirmed by selective enzymatic degradation by the M213R *Rg*DAAO variant (4 h, 30 °C) (Sacchi et al., 2002). Amino acids concentrations are expressed as nmol to mg total protein content.

### 2.5 Statistical analysis

Electrophysiological data were analyzed offline with Clampfit software (Molecular Devices, Sunnyvale, CA, USA). For LTP experiments, fEPSP amplitudes were recorded for 20 min before the TBS protocol application to establish a baseline. The potentiation of fEPSP amplitude was normalized to baseline values and expressed as a percentage. Statistical significance was evaluated over the last 10 min (70-80 min or 50-60) of recording after the conditioning stimulus. Amplitude and frequency of spontaneous synaptic currents were evaluated over 3 min of recordings. Data distribution was assessed for normality using the Shapiro–Wilk test (Supplementary Table 1). Datasets were analyzed with parametric or non-parametric tests as appropriate. Electrophysiological data were analyzed using either paired or unpaired Student’s t-tests, as appropriate. Biochemical data were analyzed using GraphPad Prism 9.0 software. Statistical significance was assessed using unpaired two-tailed Student’s *t*-test or Mann-Whitney test for comparisons between WT and *Ddo*-KI samples from mice of the same age, and by two-way ANOVA followed by multiple comparison tests for age-dependent analyses. No formal statistical test for outlier detection was applied.

All data are presented as mean ± S.E.M (standard error of the mean), and statistical significance was set at p-value < 0.05. The value of *n* refers to the number of brain slices or neurons.

## 3. Results

### 3.1 Juvenile and adult Ddo-KI mice do not exhibit alterations in basal glutamatergic transmission

As an initial assessment, we investigated hippocampal glutamatergic transmission in *Ddo*-KI mice relative to WT littermates. To probe presynaptic function, we measured paired-pulse ratio (PPR), a presynaptic readout that reflects neurotransmitter release probability. Recordings were obtained from hippocampal slices prepared from both male and female P30 and P60 mice. At both ages, *Ddo*-KI mice displayed PPR values indistinguishable from WT littermates across all interpulse intervals tested (Fig. 1A,B), indicating that presynaptic release probability is not altered by constitutive D-Asp depletion. Given prior reports of significant sex-dependent effects on glutamatergic transmission (Locklear et al., 2016; reviewed in Giacometti et al., 2020; Knouse et al., 2022), we also analyzed males and females separately at both ages. However, no significant genotype-dependent differences were observed across ages in either sex (Fig. 1C-F).

**Figure 1.**
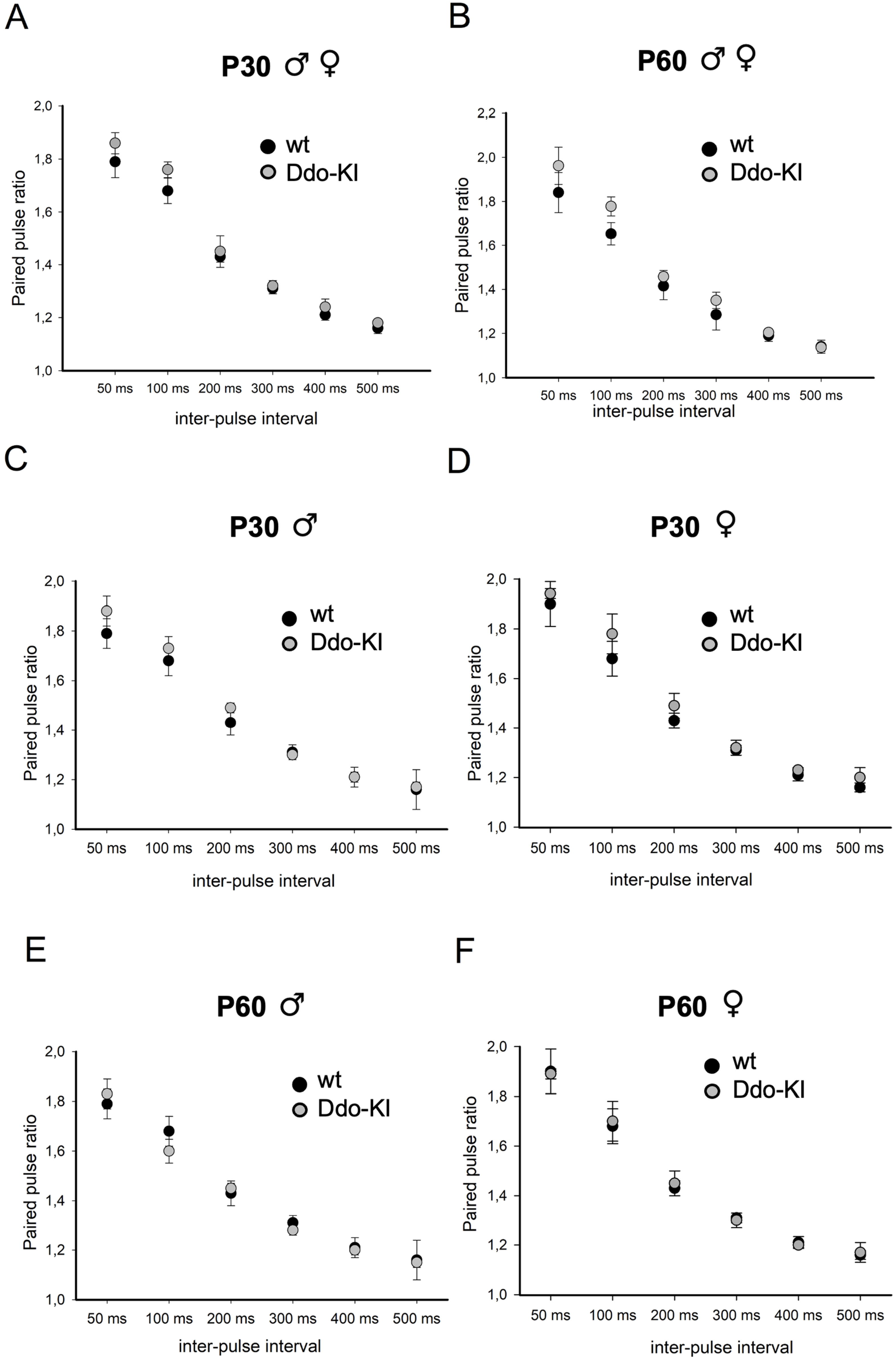
*Ddo*-KI mice do not show significant alterations in basal neurotransmission compared to wild-type littermates. Paired pulse ratio (PPR) curves plotted as fEPSP2/fEPSP1 across inter-stimulus intervals (50, 100, 200, 300, 500 ms). No statistically significant differences are observed (n = 6 males, 6 females for each genotype and age). Data are expressed as the mean PPR ± S.E.M across inter-pulse intervals. Statistical analysis was performed using an unpaired two-tailed Student’s t-test.

### 3.2 Juvenile Ddo-KI mice exhibit alterations in NMDAR-mediated hippocampal synaptic plasticity

Previous behavioral studies reported improved cognitive performance in *Ddo*-KI mice relative to WT controls (De Rosa et al., 2020). To determine whether D-Asp depletion affects the physiological mechanism supporting learning and memory, we examined NMDAR-dependent LTP in juvenile and adult mice, initially pooling males and females. At P30, *Ddo*-KI mice exhibited significantly enhanced hippocampal LTP compared with WT littermates (p= 1.168e-17, two-tailed Student’s t test, Fig. 2A), indicating that early postnatal D-Asp depletion strengthens plasticity mechanisms. This enhancement was not observed at P60 (p=0.392, two-tailed Student’s t test, Fig. 2B), suggesting a developmentally restricted phenotype.

**Figure 2.**
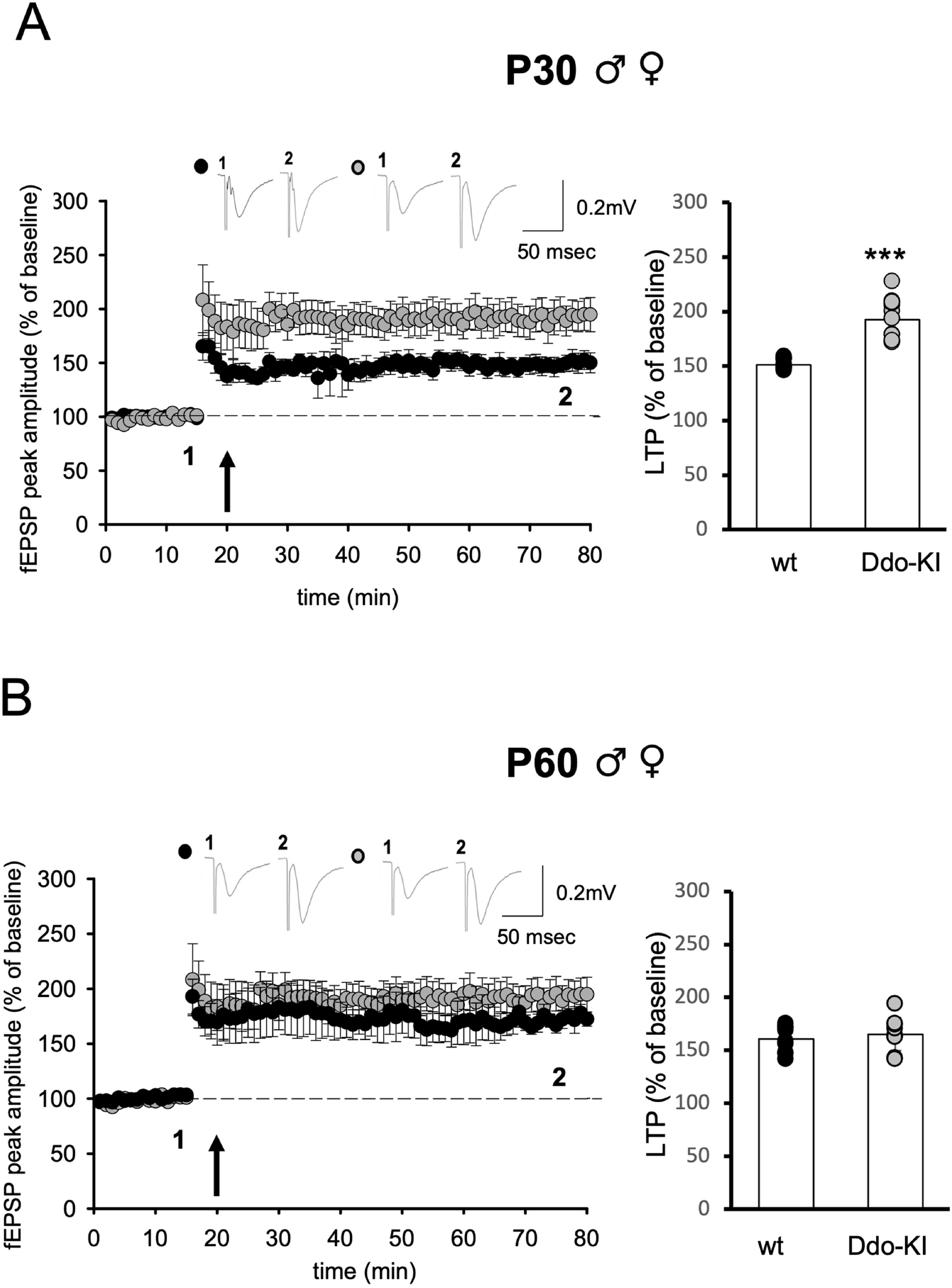
Juvenile (P30) but not adult (P60) *Ddo*-KI mice display altered hippocampal synaptic plasticity. Representative traces for each genotype and experimental condition are shown on the left; below, superimposed pooled recordings illustrate theta-burst stimulation-induced LTP in hippocampal slices from P30 (A) and P60 (B) wild-type (WT) (P30: n = 12; P60: n = 10) and *Ddo*-KI mice (P30: n = 14; P60: n = 10). Bar graphs on the right show LTP magnitude (expressed as % of baseline) for each genotype. Statistical analysis was performed using an unpaired two-tailed Student’s t-test.

When males and females were analyzed separately to assess sex-dependent effects, we found that the LTP enhancement observed in P30 *Ddo*-KI mice was specific to males relative to sex-matched WT controls (p=1.14e-13, two-tailed Student’s t test, Fig. 3A), whereas *Ddo*-KI females displayed LTP magnitudes comparable to controls (p=0.419, two-tailed Student’s t test, Fig. 3B). However, a closer examination of the induction phase revealed that female *Ddo*-KI mice exhibited a transient but significant enhancement of synaptic plasticity within 30 min after theta-burst stimulation (TBS) (Collingridge 2025) (Fig. 3C), indicating an effect restricted to the early phase of LTP. These results point to a sex-specific sensitivity to early D-Asp depletion, with male *Ddo*-KI mice showing a robust strengthening of NMDAR-dependent plasticity, as further supported by direct comparison between *Ddo*-KI males and females (p=3.412e-12, two-tailed Student’s t test, Fig. 3D).

**Figure 3.**
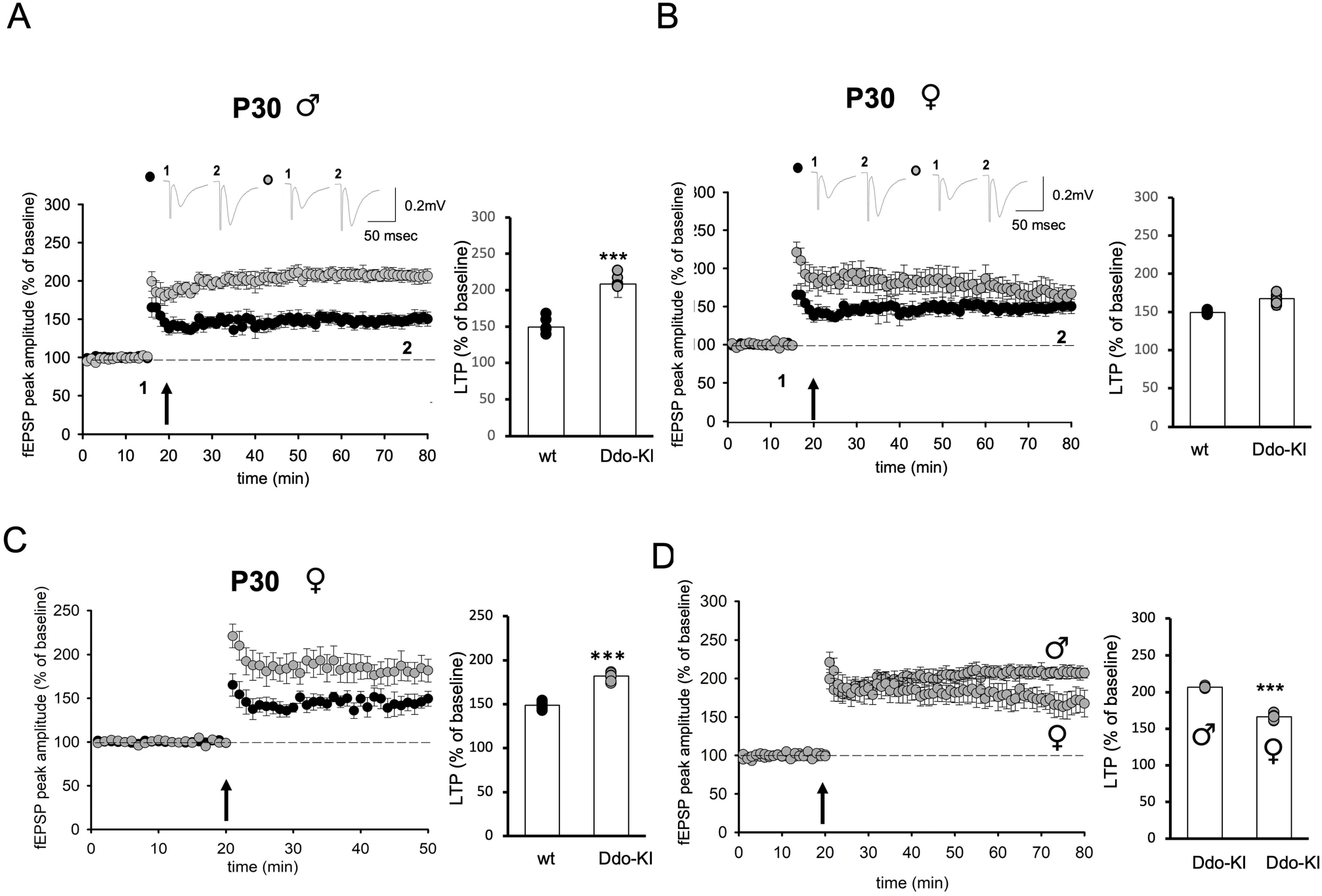
Juvenile *Ddo*-KI mice display hippocampal sex-dependent alterations in NMDA-mediated synaptic plasticity compared to wild-type littermates. (A) Representative traces for each genotype and experimental condition are shown on the left; below, superimposed pooled recordings illustrate theta-burst stimulation-induced LTP in hippocampal slices from juvenile male (P30) wild-type (WT) (n = 6) and *Ddo*-KI mice (n =7). Bar graphs on the right show LTP magnitude (expressed as % of baseline) for each genotype. (B) Representative traces for each genotype and experimental condition are shown on the left; below, superimposed pooled recordings illustrate theta-burst stimulation-induced LTP in hippocampal slices from female (P30) wild-type (WT) (n = 6) and *Ddo*-KI mice (n =7). Bar graphs on the right show LTP magnitude (expressed as % of baseline) for each genotype. (C) Superimposed pooled recordings illustrate the LTP induction phase, 40 minutes after theta-burst stimulation-induced LTP in hippocampal slices from females (P30). Bar graphs on the right show LTP magnitudes (expressed as % of baseline) for each genotype. (D) Superimposed pooled recordings illustrate theta-burst stimulation-induced LTP in hippocampal slices from male (P30) and female (n=7) *Ddo*-KI mice. Bar graphs on the right show LTP magnitudes (expressed as % of baseline) for each sex. Statistical analysis was performed using an unpaired Student’s t-test.

Given the LTP enhancement observed in P30 *Ddo*-KI male mice, we next evaluated whether this sex-dependent effect persisted in adulthood by analyzing P60 males and females separately. In contrast to juvenile males, LTP induction in adult *Ddo*-KI male mice did not differ from that of sex-matched controls (Fig. 4A), confirming that the effect was developmentally restricted. Likewise, *Ddo*-KI females displayed LTP amplitude comparable to their respective controls (Fig. 4B). Together, these findings indicate that the enhancement of hippocampal NMDAR-dependent plasticity driven by D-Asp depletion is confined to the juvenile stage.

**Figure 4.**
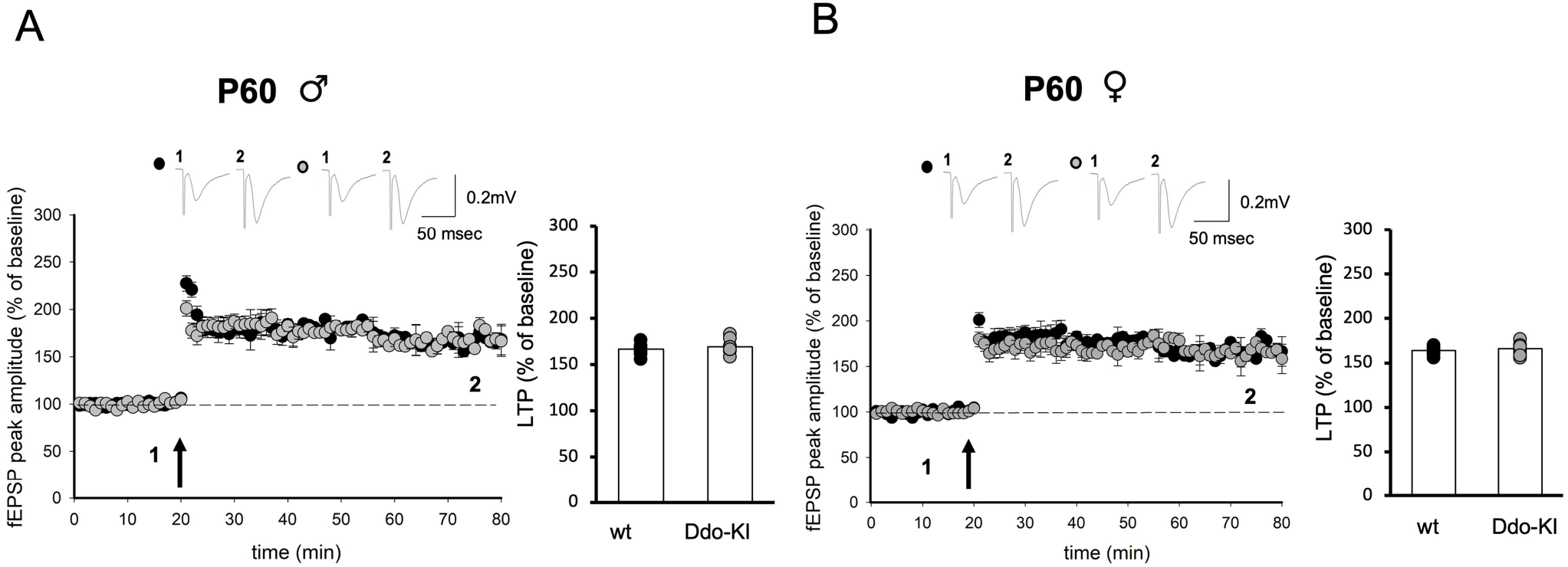
Adult *Ddo*-KI mice do not display hippocampal sex-dependent alterations. Representative traces for each genotype and experimental condition are shown on the left; below, superimposed pooled recordings illustrate theta-burst stimulation-induced LTP in hippocampal slices from adult male (P60) (A) and female (B) wild-type (WT) (n = 5) and *Ddo*-KI mice (n =5). Bar graphs on the right show LTP magnitude (expressed as % of baseline) for each genotype. Statistical analysis was performed using an unpaired Student’s t-test.

The specificity of the LTP modulation with respect to genotype, age, and sex was further explored through additional cross-comparisons between males and females of the same age and genotype, as well as between juvenile and adult mice within each sex and genotype (Supplementary Fig. 1A-F). These analyses did not reveal significant differences, supporting the conclusion that the observed phenotype is restricted to the specific experimental conditions described above.

### 3.3 D-Asp depletion during embryogenesis does not alter neurotransmitter release but affects synaptic AMPAR/NMDAR balance in P30 male Ddo-KI mice

To identify the synaptic mechanisms underlying the enhanced LTP observed in P30 *Ddo*-KI male mice, we performed whole-cell patch-clamp recordings from single hippocampal CA1 pyramidal neurons. Spontaneous excitatory and inhibitory events were measured to assess whether D-Asp depletion induces an imbalance in excitatory/inhibitory synaptic activity, characteristic of different ASD disease models (Nelson et al., 2015; De Jaco et al., 2017). Both sEPSC and sIPSC, in terms of frequency and amplitude, were unchanged between *Ddo*-KI mice and controls (Fig. 5A,B), and the E/I ratio (Fig. 5C). Presynaptic function was also unaffected, as shown by unaltered PPR values (Fig. 5D). These data indicate that the strengthened LTP in juvenile *Ddo*-KI male mice does not arise from changes in spontaneous or evoked neurotransmitter release.

**Figure 5.**
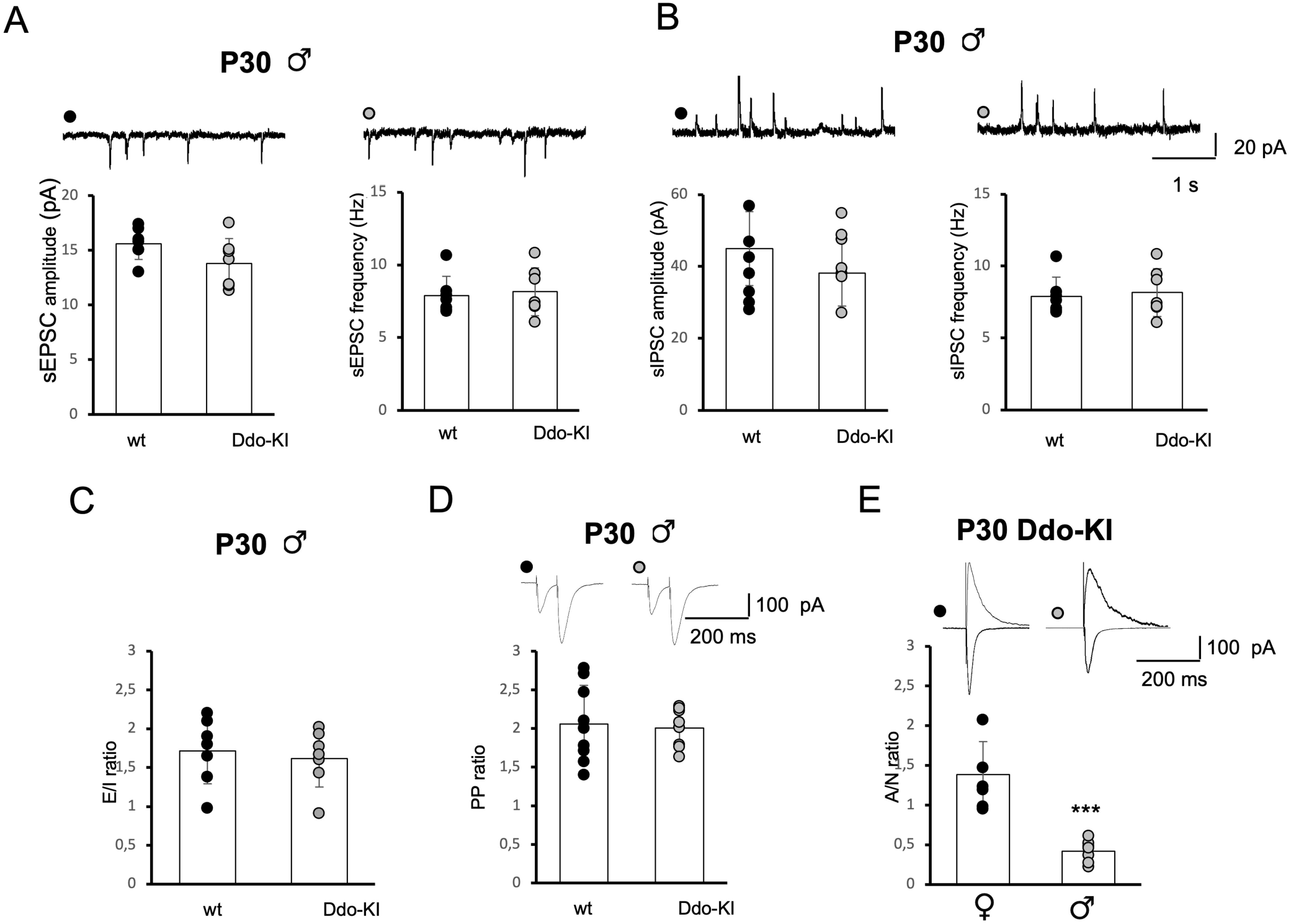
Prenatal D-Asp deficit does not affect neurotransmitter release but reduces AMPAR/NMDAR synaptic contribution in P30 male *Ddo*-KI mice compared to wild type. (A, B) Representative current traces are shown in the upper part of the panels; bar graphs show sEPSC (A) and sIPSC (B) amplitude (left) and frequency (right) in male (P30) wild-type (WT) and *Ddo*-KI mice. sEPSC: n = 7/genotype; sIPSC: n = 7/genotype. (C) Excitation/Inhibition (E/I) ratio (sEPSC frequency /sIPSC frequency) in male (P30) WT (n = 7) and *Ddo*-KI (n = 7) mice. (D) Representative current traces (top) and histogram of paired pulse (PP) ratio in male WT (n = 9) and *Ddo*-KI (n = 8) mice. (E) Representative current traces (top) and AMPAR/NMDAR ratio (bottom) in *Ddo*-KI male (n = 6) and females (n = 7). The AMPA/NMDA ratio is significantly reduced in *Ddo*-Ki male mice compared to females. Statistical analysis was performed using unpaired Student’s t-test as appropriate.

Given the absence of presynaptic or E/I alterations, we next analyzed whether the observed LTP phenotype could reflect changes in postsynaptic AMPAR and/or NMDAR current expression. Accordingly, P30 male *Ddo*-KI mice displayed a significant reduction in the AMPAR/NMDAR ratio compared with age-matched controls (p=2.08e-4, two-tailed Student’s t test, Fig. 5E), suggesting a shift in postsynaptic receptor balance as an adaptive response to D-Asp depletion.

### 3.4 D-Asp perfusion restores LTP magnitude in P30 male Ddo-KI mouse hippocampal slices

Pharmacological studies have demonstrated that the absence of an endogenous agonist can trigger compensatory receptor adjustments, typically involving a reduction in receptor number combined with increased receptor sensitivity, a phenomenon often referred to as sensitization (Charlton, 2009; Mango et al., 2014). Within this framework, we hypothesized that brief D-Asp exposure in hippocampal slices from P30 *Ddo*-KI male mice could counteract the observed sensitized state. Consistent with this prediction, acute D-Asp perfusion (2 μM, 30 min), a concentration that does not affect basal neurotransmission (Fig. 6A) or LTP (Fig. 6B) in the hippocampus of P30 WT male mice, restored LTP amplitude in *Ddo*-KI hippocampal slices to control levels (p=4.11e-24, two-tailed Student’s t test, Fig. 6C). This rescue supports the view that altered receptor balance at glutamatergic synapses in D-Asp-deficient mice reflects a reversible adaptive state rather than a structural synaptic deficit.

**Figure 6.**
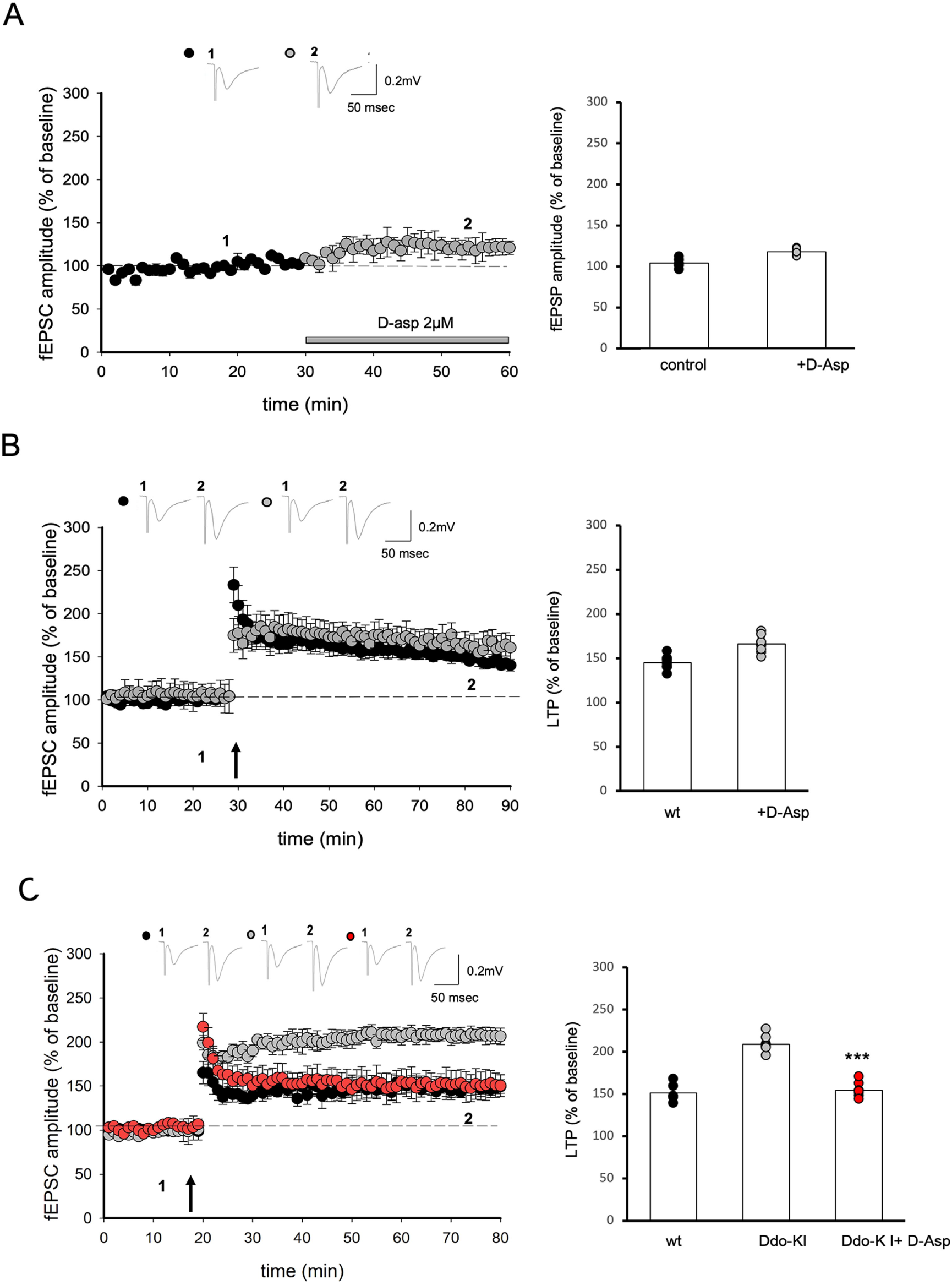
D-Asp perfusion rescues synaptic plasticity in hippocampal slices from juvenile male *Ddo*-KI mice. (A) Representative traces in control condition (1) and during 2 μM D-Asp (2) perfusion are shown on the left; below, superimposed pooled data show basal neurotransmission expressed as % fEPSP relative to control (control n = 8; D-Asp, n = 8). The bar graph on the right shows fEPSC amplitude in different conditions. (B) Representative traces for wild-type (WT) and D-Asp perfused group are shown on the left; below, superimposed pooled data show LTP values induced by theta-burst stimulation and expressed as % of baseline (WT: n = 8; D-Asp: n = 10). The bar graph on the right shows LTP magnitude (% of baseline) for experimental groups. (C) Representative traces for each experimental group are shown on the top; below, superimposed pooled recordings show theta-burst stimulation-induced LTP in slices from P30 male WT (n = 6), *Ddo*-KI (n = 8), and *Ddo*-KI perfused with D-Asp (n = 10). (B) Bar graphs show LTP magnitude (expressed as % of baseline). D-Asp application significantly reduces LTP amplitude in *Ddo*-KI mouse hippocampal slices, restoring values comparable to WT mice. Statistical analysis was performed using paired and unpaired Student’s *t*-test as appropriate.

### 3.5 DASPO activity and D-Asp levels in Ddo-KI hippocampi do not change across ages

To determine whether the transient LTP alterations observed between juvenile and adult *Ddo*-KI mice could be attributable to enhanced D-Asp catabolism, we measured DASPO enzymatic activity and D-Asp levels in hippocampal samples from P30 and P60 mice. DASPO activity was approximately 2–3-fold higher in *Ddo*-KI mice than in WT at both ages, reaching statistical significance at P60 (p = 0.0040, two-tailed Student’s t test), and showing a similar trend at P30, although not reaching statistical significance (p = 0.0622) (Fig. 7A). Consistently, age dependent-analysis by two-way ANOVA revealed a significant genotype effect [F_(1, 13)_ = 18.06, p = 0.0009] but no significant effect of age [F_(1, 13)_ = 1.193, p = 0.2945] nor genotype x age interaction [F_(1, 13)_ = 0.4519, p = 0.5132]. Tukey’s post-hoc test revealed no significant differences between P30 and P60 within *Ddo*-KI mice (p = 0.9897), while confirming a significant increase in *Ddo*-KI mice compared to WT at P60 (p = 0.0219) (Fig. 7A). As expected, D-Asp levels were significantly lower in *Ddo*-KI hippocampi compared with WT, at both P30 (p = 0.0320, two-tailed Student’s t test) and P60 (p = 0.0078) (Fig. 7B), while L-Asp levels were unchanged between *Ddo*-KI and WT mice (Fig. 7B). As a result, the D-Asp/total Asp ratio was significantly reduced at both P30 (p = 0.0382) and P60 (p = 0.0135). The levels of other relevant amino acids, including D-Ser, L-Ser, Gly, L-Asn, L-Gln, and L-Glu, were not significantly affected by DASPO overexpression across ages in *Ddo*-KI mice (Supplementary Fig. 2).

**Figure 7.**
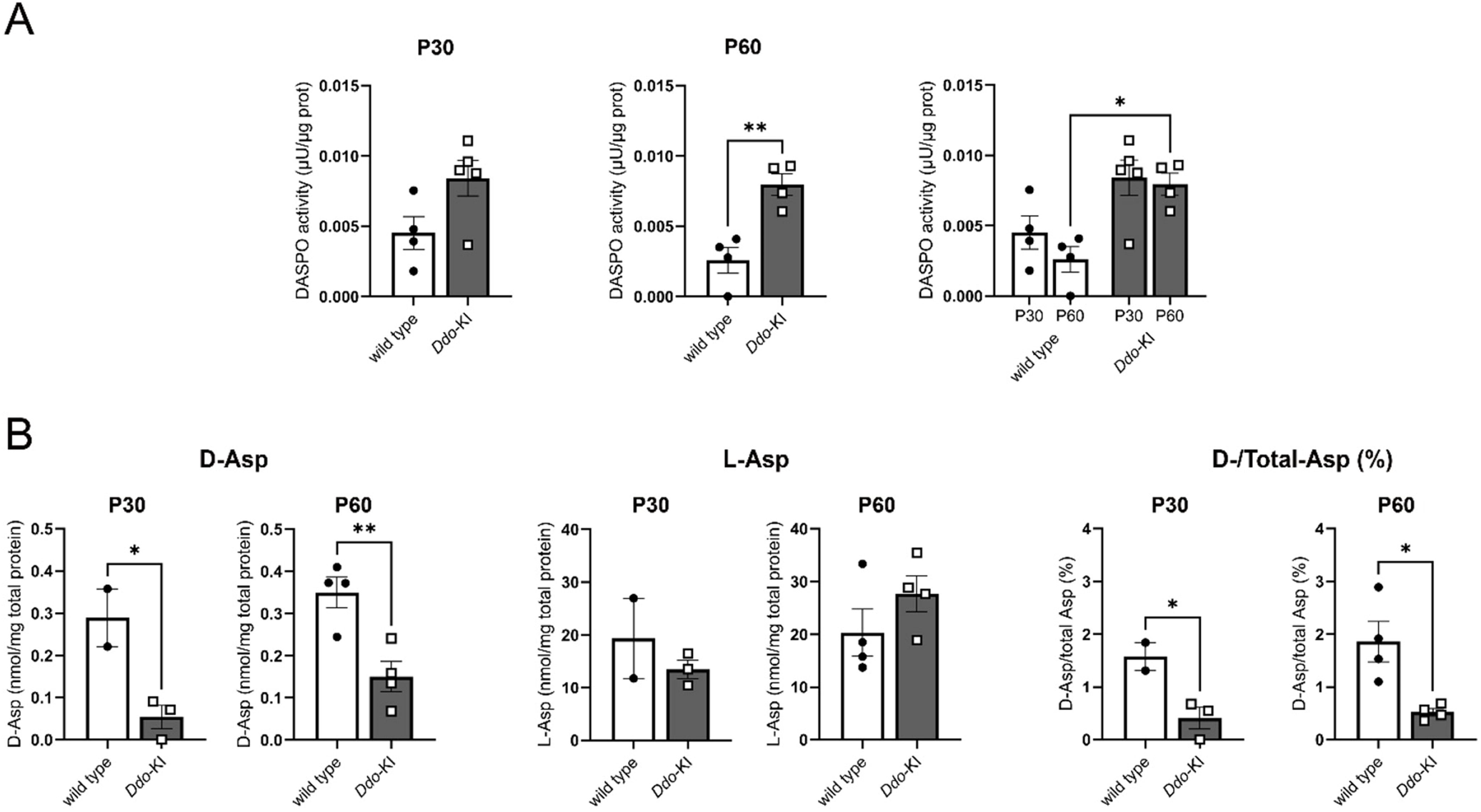
Persistent DASPO enzymatic activity increase and D-Asp depletion across developmental stages in the hippocampus of *Ddo*-KI mice. DASPO enzymatic activity (A), D-Asp, L-Asp levels and D-Asp/total Asp (%) (B) in P30 and P60 Ddo-KI and wild-type (WT) mice. Data are presented as mean ± SEM with individual data points (dots), each representing one biological sample analyzed in at least two technical replicates. DASPO enzymatic activity was expressed as µU/µg protein, amino acid levels as nmol/mg total protein, and D-Asp/total Aspo ratio as percentage (%). Statistical analyses were performed using unpaired two-tailed Student’s *t*-test for comparisons between WT and *Ddo*-KI samples from mice of the same age, and by two-way ANOVA for age-dependent analyses, followed by Tukey’s post-hoc analysis.

Collectively, these findings indicate that DASPO overexpression and the resulting D-Asp downregulation remain stable between juvenile and adult stages, and therefore are unlikely to account for the LTP differences observed.

## 4. Discussion

A hallmark of D-Asp physiology is its tightly regulated developmental trajectory, with levels peaking during embryonic and early postnatal stages and declining sharply after birth (Dunlop et al., 1986; Neidle and Dunlop, 1990; Wolosker et al., 2000), due to postnatal DASPO upregulation (Hashimoto et al., 1993,1995; Sakai et al., 1998; Errico et al., 2018; De Rosa et al., 2020; Punzo et al, J Neurosci, 2016). This profile suggests that D-Asp acts as a transient regulator of neurodevelopmental processes, including neurogenesis, axon outgrowth, synaptogenesis, and circuit maturation. Consistently, D-Asp is enriched in proliferative and migratory zones during early development and later becomes concentrated at synaptic terminals (Sakai et al., 1998), further supporting the notion that perturbations in D-Asp homeostasis during critical developmental windows may exert enduring consequences on brain maturation and function.

In the present study, we show that constitutive D-Asp depletion in *Ddo*-KI mice leads to a paradoxical increase in NMDAR-dependent LTP at CA1 synapses during the juvenile stage (P30). This finding is particularly intriguing in light of previous evidence demonstrating that increased D-Asp availability, such as in *Ddo* knockout or D-Asp-treated mice, also enhances hippocampal LTP and cognitive performance (Errico et al., 2008; Errico et al., 2008; Errico et al., 2011a,b; Errico et al., 2014).

One plausible interpretation is that increased D-Asp levels in adult mice exerts a direct facilitatory effect on NMDAR activation, thereby enhancing synaptic plasticity under physiological or elevated levels. In contrast, embryonic D-Asp depletion may trigger compensatory homeostatic mechanisms, leading to increased glutamatergic receptor sensitivity or altered receptor composition, which align with recent proteomic findings reported in the brain of *Ddo*-KI mice at P3 (Errico et al., 2025). In this scenario, the enhanced LTP observed in *Ddo*-KI mice would not reflect improved synaptic function per se, but rather a state of maladaptive plasticity arising from chronic neuromodulator deficiency. Accordingly, the absence of an endogenous NMDA receptor ligand would be expected to weaken, rather than potentiate, hippocampal synaptic plasticity.

Importantly, this effect was limited to a defined temporal window, as LTP normalized by adulthood (P60), indicating that the impact of D-Asp deficiency is strongly dependent on developmental context. Interestingly, DASPO activity remained persistently sustained and D-Asp levels persistently reduced across ages, suggesting that the enhanced synaptic plasticity does not arise from progressive metabolic changes but rather from a transient phase of heightened circuit sensitivity. We propose that during early stages, when hippocampal circuits are highly plastic and more dependent on D-Asp signaling (P30), its absence triggers compensatory adjustments that enhance NMDAR-related synaptic responsiveness. As maturation progresses and circuit plasticity declines (P60), these adaptive processes stabilize, reducing the functional impact of D-Asp deficiency. However, identifying the precise mechanisms underlying this feedback upregulation of LTP in juvenile mice is challenging, as homeostatic upscaling and downscaling processes regulating synaptic plasticity are highly complex and vary significantly across brain regions, neuronal types, and even individual synapses (Pozo and Goda, 2010).

Mechanistically, our data indicate that this phenotype is primarily postsynaptic. Indeed, basal synaptic transmission, short-term presynaptic plasticity and excitatory/inhibitory balance were all preserved in *Ddo*-KI hippocampal circuits. Conversely, we detected a reduced AMPAR/NMDAR ratio in juvenile P30 *Ddo*-KI mice, consistent with a shift in postsynaptic receptor balance. This finding is consistent with a model of network reorganization aimed at maintaining glutamatergic synaptic function or composition in the absence of proper physiological D-Asp signaling. Supporting this interpretation, our previous proteomic findings in early postnatal (P3) *Ddo*-KI brains showed widespread changes in synaptic proteins, including the NMDAR subunit GluN1, the AMPAR subunits GluA3 and GluA4, as well as the postsynaptic density proteins Homer1 and Homer3, together with additional molecules directly or indirectly interacting with NMDARs and/or mGluR5 (Errico et al., 2025).

Importantly, the enhanced LTP observed in juvenile *Ddo*-KI mice was rapidly normalized by acute bath application of exogenous D-Asp, indicating that the phenotype is reversible and does not reflect permanent structural abnormalities in the circuit. Rather, these findings support the existence of a dynamic compensatory state in which synaptic function is continuously tuned in response to D-Asp availability.

A further layer of complexity is introduced by the sex-dependent component of LTP since we observed that LTP enhancement in juvenile *Ddo*-KI mice was more pronounced in males during the consolidation phase, whereas females exhibited a transient increase restricted to the induction phase. This divergence suggests that D-Asp depletion interacts with sex-specific developmental trajectories of glutamatergic signaling, in line with previous evidence indicating sex-dependent regulation of NMDAR function and synaptic plasticity (Monfort et al., 2015; Knouse et al., 2022; Kniffin et al., 2024). These differences may reflect variations in receptor composition, intracellular signaling pathways, or hormonal modulation and warrant further investigation to elucidate their underlying mechanisms.

Destabilization of neural circuits by excessive potentiation has been associated with neurodevelopmental disorders and proposed as a mechanism contributing to ASD-related phenotypes (Chen *et al*. 2024). In this context, both clinical and preclinical evidence links altered D-Asp metabolism to neuropsychiatric phenotypes. Reduced D-Asp levels and increased DDO activity have been reported in schizophrenia brains (Errico et al., 2013; Nuzzo et al., 2017), while altered D-Asp regulation has also been described in ASD-related models (Nuzzo et al., 2020; De Rosa et al., 2022; Di Maio et al., 2025). Moreover, *Ddo*-KI mice display behavioral alterations, including enhanced memory and impaired sociability (De Rosa et al., 2020; Lombardo et al., 2022), and a rare human case of *DDO* duplication has been associated with neurodevelopmental abnormalities (Lombardo et al., 2022). Together, these findings suggest that dysregulation of D-Asp-dependent plasticity may contribute to circuit-level dysfunction despite apparent enhancements in synaptic potentiation. Thus, the apparent gain in synaptic strength observed in *Ddo*-KI mice may reflect compensatory overactivation rather than functional improvement.

In summary, this study identifies embryonic D-Asp as a critical modulator of NMDAR-dependent synaptic plasticity during hippocampal development. Using a model of constitutive D-Asp depletion, we revealed a transient, juvenile-specific enhancement of LTP driven by postsynaptic compensatory mechanisms and reversible upon D-Asp reintroduction. These findings highlight a previously unrecognized role of D-Asp in shaping developmental trajectories of hippocampal synaptic function and underscore its potential implication in neuropsychiatric vulnerability associated with glutamatergic dysfunction.

## Abbreviations

AMPAR: α-amino-3-hydroxy-5-methyl-4-isoxazolepropionic acid receptors
ASD: autism spectrum disorders
CNS: central nervous system
DDO: D-aspartate oxidase
D-Asp: D-Aspartate
EPSC: excitatory postsynaptic currents
fEPSP: field excitatory post-synaptic potential
sEPSC: spontaneous excitatory postsynaptic currents
sIPSC: spontaneous inhibitory postsynaptic currents
KI: knock-in
LTP: long-term potentiation
mGluR5: metabotropic glutamate receptor 5
NMDAR: N-methyl-D-aspartate receptors
PP: paired pulse
PPR: paired pulse ratio
PFC: prefrontal cortex
TBS: theta burst stimulation

## Author Contributions

**DM**: investigation, data curation, formal analysis, writing– original draft, writing – review and editing; **FE:** conceptualization, data curation, formal analysis, writing – review and editing, writing – original draft; **ZM**: investigation, data curation, formal analysis; **SD**: investigation, data curation, formal analysis; **ADM**: investigation, data curation, formal analysis; **RN:** conceptualization; **LP**: investigation, data curation, formal analysis; writing, review and editing; **MEDS**: conceptualization, writing, review and editing; **AU**: conceptualization, data curation, formal analysis, writing – review and editing, writing – original draft.

## Data Availability Statement

The data that support the findings of this study are available from the corresponding author upon reasonable request.

