## Supplemental material for "Embryonic depletion of D-aspartate perturbs NMDA receptor-dependent long-term potentiation in the hippocampus of juvenile mice"

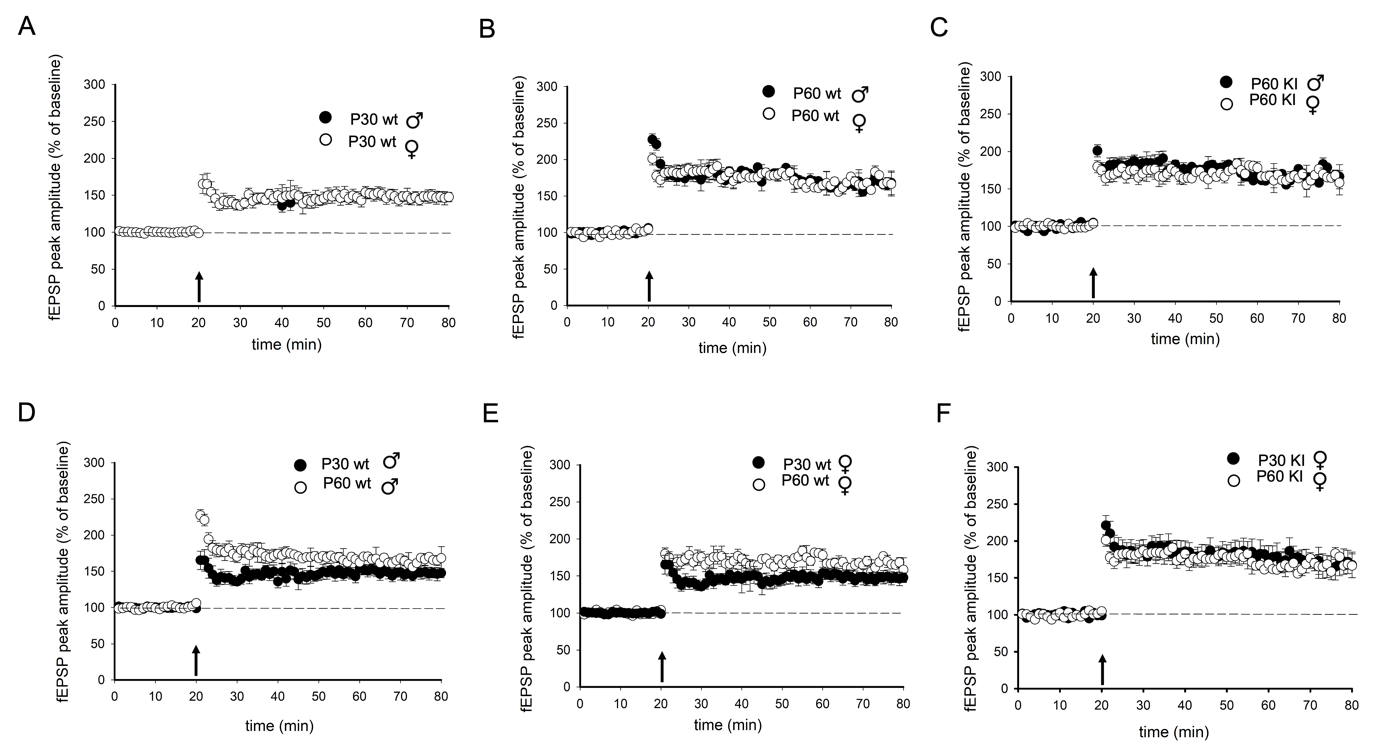

**Supplementary Figure 1. Age- and sex-dependent comparison of synaptic plasticity in wild type and Ddo-KI mice** (A-C) Representative superimposed pooled recordings illustrating theta-burst stimulation-induced LTP in hippocampal slices from P30 (A) and P60 (B,C) male and female wild type (wt) (A,B) and Ddo-KI (C) mice. (D-F) Representative superimposed pooled recordings illustrating theta-burst stimulation-induced LTP in hippocampal slices from male (D) and female (E,F) wt (D,E) and Ddo-KI (F) mice. No significant differences are detected in any of the comparisons. Statistical analyses were performed using Student’s t-test, based on data distribution assessed by the Shapiro–Wilk normality test.

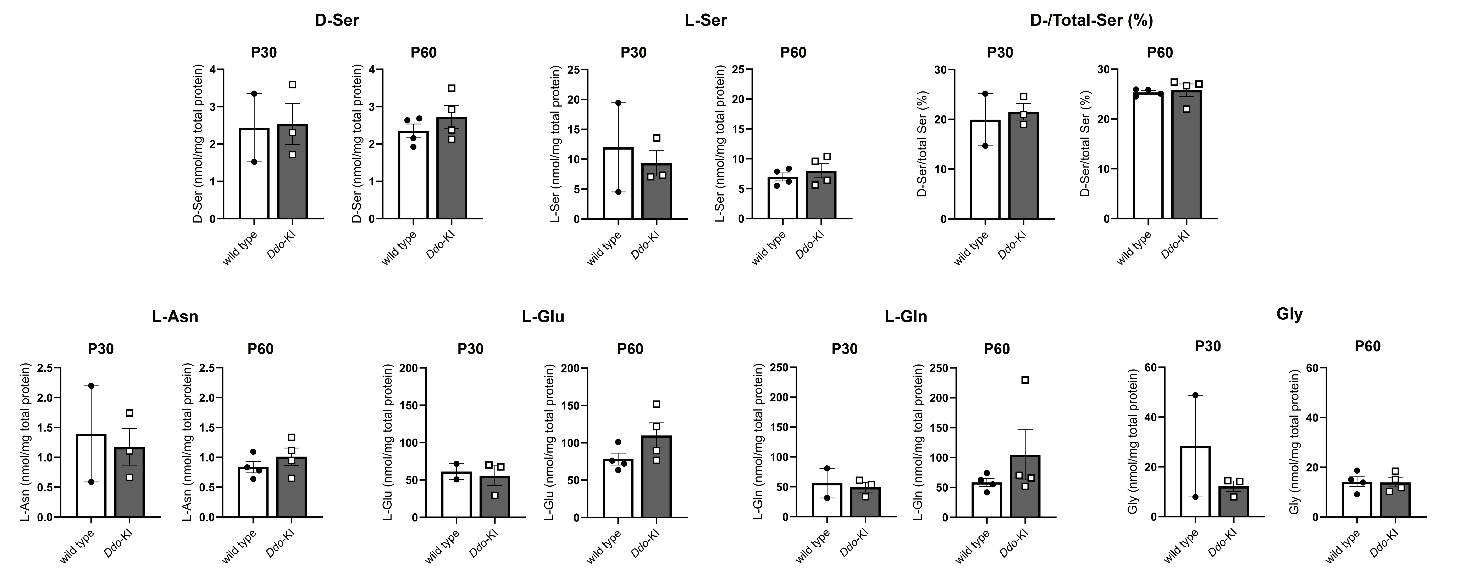

**Supplementary Figure 2.** **Amino acid levels (nmol/mg total protein) and D-Ser/total Ser (%) in P30 and P60 wild-type and *Ddo*-KI mice.** Data are presented as mean ± SEM with individual data points (dots), each representing one biological replicate analyzed in at least three technical replicates. Statistical analyses were performed using Student’s t-test or the Mann–Whitney test, as appropriate, based on data distribution assessed by the Shapiro–Wilk normality test. The Mann–Whitney test was applied to D-Ser/total Ser ratio and L-glutamine, while all other analytes were analyzed using Student’s t-test.

**Supplementary Table 1.** Results of the normality tests.

|  | **p value (Shapiro-Wilk test)** | | | | |
| --- | --- | --- | --- | --- | --- |
|  | **P30** | |  | **P60** | |
|  | **Wt** | **DDO KI** |  | **Wt** | **DDO KI** |
| **D-Asp** | N.A. | 0,389 |  | 0,154 | 0,939 |
| **L-Asp** | N.A. | 0,974 |  | 0,140 | 0,808 |
| **D-Asp Ratio** | N.A. | 0,351 |  | 0,766 | 0,962 |
| **D-Ser** | N.A. | 0,599 |  | 0,341 | 0,752 |
| **L-Ser** | N.A. | 0,064 |  | 0,484 | 0,340 |
| **D-Ser Ratio** | N.A. | 0,683 |  | 0,361 | **0,030** |
| **L-Asn** | N.A. | 0,800 |  | 0,724 | 0,976 |
| **L-Glu** | N.A. | 0,104 |  | 0,418 | 0,727 |
| **L-Gln** | N.A. | 0,474 |  | 0,994 | **0,018** |
| **Gly** | N.A. | 0,101 |  | 0,888 | 0,767 |
| **DASPO** | 0,9348 | 0,152 |  | 0,315 | 0,396 |
| **PPR ♂♀** | 0,915 | 0,893 |  | 0,876 | 0,935 |
| **PPR ♂** | 0,905 | 0,912 |  | 0,921 | 0,987 |
| **PPR ♀** | 0,956 | 0,924 |  | 0,92 | 0,923 |
| **LTP ♂♀** | 0,745 | 0,649 |  | 0,78 | 0,736 |
| **LTP ♂** | 0,748 | 0,799 |  | 0,738 | 0,807 |
| **LTP ♀** | 0,75 | 0,823 |  | 0,793 | 0,836 |
| **A/N Ratio ♂** | - | 0,961 |  | - | - |
| **A/N Ratio♀** | - | 0,901 |  | - | - |
| **PP Ratio ♂** | 0,933 | 0,766 |  | - | - |
| **sEPSC ♂** | 0,857 | 0,895 |  | - | - |
| **sIPSC ♂** | 0,906 | 0,788 |  | - | - |
| **E/I Ratio ♂** | 0,767 | 0,959 |  | - | - |
| **LTP +D-Asp** | 0,864 | - |  | - | - |
| **LTP +D-Asp** | 0,743 | 0,813 |  | - | - |

Abbreviations: N.A.: Not Applicable; -: group was non-used for this specific protocol; D-Asp: D-Aspartate; L-Asp: L-Aspartate; D-Ser: D-Serine; L-Ser: L-Serine; L-Asn: L-Asparagine; L-Glu: L-Glutamate; L-Gln: L-Glutamine, L-Gly: L-Glycine, sIPSC: spontaneous Inhibitor Post Synaptic Current; sEPSC: spontaneous Excitatory Post synaptic current; PR: Paired Pulse Ratio; E/I: Excitatory/inhibitory Ratio; A/N: AMPAR/NMDAR Ratio; LTP: Long Term Potentiation; DASPO: D-aspartate oxidase; Wt: Wilde type; DDO-KI: Ddo-knock-in mice.
